# DNA Methylation Landscape Reflects the Spatial Organization of Chromatin in Different Cells

**DOI:** 10.1101/080531

**Authors:** Ling Zhang, Wen Jun Xie, Sirui Liu, Luming Meng, Chan Gu, Yi Qin Gao

## Abstract

The relation between DNA methylation and chromatin structure is still largely unknown. By analyzing a large set of sequencing data, we observed a long-range power law correlation of DNA methylation with cell-class-specific scaling exponents in the range of thousands to millions of base pairs. We showed such cell-class-specific scaling exponents are caused by different patchiness of DNA methylation in different cells. By modeling the chromatin structure using Hi-C data and mapping the methylation level onto the modeled structure, we demonstrated the patchiness of DNA methylation is related to chromatin structure. The scaling exponents of the power law correlation is thus a display of the spatial organization of chromatin. Besides, the local correlation of DNA methylation is associated with nucleosome positioning and different between partially-methylated-domain and non-partially-methylated-domain, suggesting their different chromatin structures at several nucleosomes level. Our study provides a novel view of the spatial organization of chromatin structure from a perspective of DNA methylation, in which both long-range and local correlations of DNA methylation along the genome reflect the spatial organization of chromatin.

## Introduction

Composed of DNA and histone proteins, chromatin has a three-dimensional (3D) structure at different hierarchical levels (Gibcus & Dekker, 2013). The spatial organization of chromatin plays an essential role in many genomic functions, including gene expression, DNA replication and cell mitosis (Dekker, 2008; Galupa & Heard, 2015; Levine et al, 2014; Naumova et al, 2013; Pombo & Dillon, 2015). Several lines of evidence show that epigenetics can remodel chromatin structure at different levels (Aranda et al, 2015; Bell & Felsenfeld, 2000; Boettiger et al, 2016; Cedar & Bergman, 2009; Chodavarapu et al, 2010; Jenuwein & Allis, 2001). Super-resolution imaging recently showed that chromatin folding varies for different epigenetic states (Boettiger et al, 2016).

DNA methylation, as the most abundant epigenetic modification in eukaryotic chromosomes, is also thought to influence chromatin structure (Cedar & Bergman, 2009). DNA methylation has a close relationship with nucleosome positioning (Chodavarapu et al, 2010), and the binding of CTCF can be partly influenced by DNA methylation and thus changes chromatin structure (Bell & Felsenfeld, 2000). Recently, DNA methylation was also used to reconstruct A/B compartments of chromatin revealed by Hi-C experiments (Fortin & Hansen, 2015). Nevertheless, how DNA methylation relates with chromatin structure remains largely unknown.

On the other hand, the distribution of DNA methylation in chromatin, and thus the correlation of DNA methylation levels between different genomic segments, may provide hints on the spatial organization of chromatin. Here we investigate the long-range and local correlations in DNA methylation landscape, which we expect to reflect the packing of DNA in the 3D space, and try to obtain information on the underlying chromatin structure. DNA methylation possesses long-range power law correlation with cell-class-specific scaling exponent. In addition, the scaling exponent can be used to discern cell classes. We find that the degree of DNA methylation patchiness is cell-specific and this patched methylation pattern contributes to the different scaling exponents in different cells. Using polymer modeling with Hi-C data, we show that the partially-methylated-domains (PMDs) spatially segregate from the non-PMDs (genomic regions that aren’t classified as PMD) in IMR90 cell line, leading to its patchiness of DNA methylation different from h1 cell line. In this way, the cell-class-specific exponents for the long-range DNA methylation correlation reflect the spatial organization of chromatin. We also demonstrate that the local DNA methylation correlation is related with chromatin structure at several nucleosomes level, and suggests the different chromatin structure of PMD and non-PMD. Therefore, both long-range and local DNA methylation correlations can reflect the spatial organization of chromatin.

## Results

### DNA methylation shows long-range power law correlation

We compared the Pearson correlation coefficients of DNA methylation levels (beta values) within the methylome across a wide-range of human cells, including normal somatic cells, cancer cells, brain cells, gland cells and stem cells. In calculating the long-range correlation, the methylation level was first averaged using a 200-bp window. The sources of relative whole-genome bisulfite sequencing data were summarized in Appendix Data (Appendix Table S1-S5).

Taking chromosome 1 as representative, all the methylation correlations strikingly present a long-range power law decay as the genomic distance increases (Fig 1). The power law correlation implies a scale-free property of DNA methylation and the scale-invariant genomic segment lies in the kilobase-to-megabase scale. Power law scaling is of general interest (Clauset et al, 2009) and often noticed in evolving systems that may be produced by hierarchical structure of several length-scales (Eugene et al, 2006). The scale-invariant genomic scale (kilobase-to-megabase) is also the sizes of genes and chromatin domains, and can be important for a variety of genomic functions (Rao et al, 2014; Schneider & Grosschedl, 2007).

**Figure 1.**
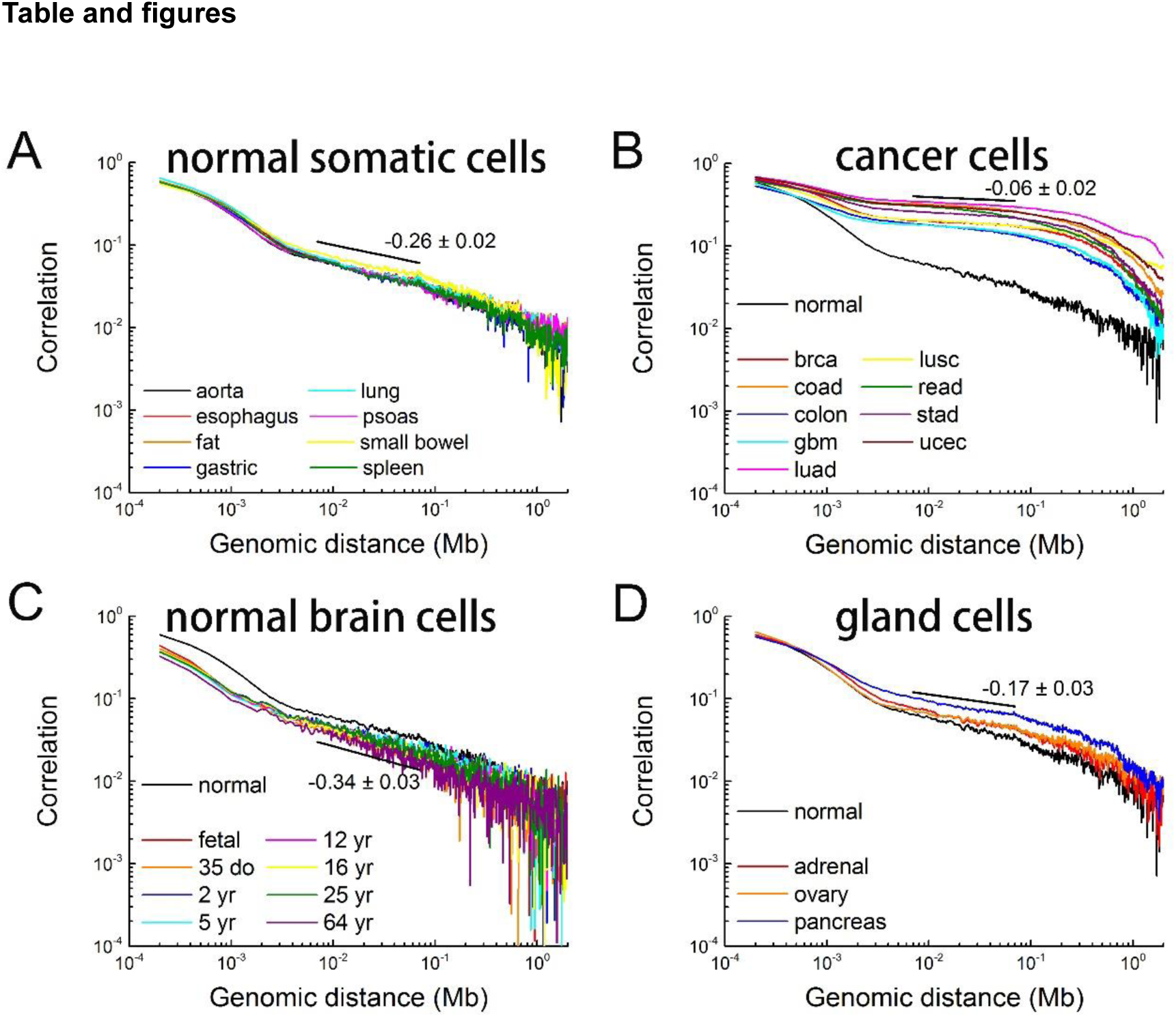
Long-range correlations in DNA methylation are distinct among different cell classes. The Pearson correlation coefficients for chromosome 1 of different cells are shown in log-log plots. The average scaling exponents are annotated in the figure. A, 8 different somatic cells: aorta, esophagus, fat, gastric, lung, psoas, small bowel and spleen. B, 9 different cancer cells: brca, coad, colon, gbm, luad, lusc, read, stad and ucec. The cells are labelled following TCGA except for colon cancer (colon). C, Normal brain cells of different ages (fetal, 35 days old, 2-64 years old). D, 3 different gland cells (adrenal, ovary, pancreas). Correlation for normal aorta cells (normal) is also plotted for comparison in B, C and D.

The correlation coefficients still have finite values in the order of 0.01-0.1 even for the 1 Mb genomic separation (Fig 1). To verify the statistical significance of the power law decay found here, we also calculated the correlation of a randomly methylated DNA sequence for comparison. Specifically, we generated a randomized methylation pattern by randomly assigning the methylation level of each CpG following the overall distribution of the original sample. The correlation coefficient for the random sample immediately drops to zero and the power law decay disappears (Appendix Fig S1). This comparison clearly shows that the non-random nature of DNA methylation in the cells and that the methylation level of CpGs separated by a very long genomic distance is indeed significantly correlated.

Interestingly, the scaling exponents of long-range DNA methylation differ substantially between normal somatic cells and cancer cells, and the respective values are −0.26±0.02 and −0.06±0.02. The value for cancer cells is significantly smaller than that for normal somatic cells. Small standard deviations show that the scaling exponents are conserved among either normal somatic cells or cancer cells (Figs 1A and 1B), although the methylation levels of individual CpGs (and even the average values among all CpGs) vary greatly (Schultz et al, 2015). Similarly, the differences between normal somatic cells and brain cells or gland cells are substantial but consistent within each cell class (Figs 1C and 1D), suggesting that cellular differentiation causes systematic variations of DNA methylation landscape. It was found that the scaling exponents for chromosome 1 of normal somatic cells in three different individuals are conserved (Fig 2A, Appendix Figs S2A and S3B) and the power law scaling is also present in mouse brain cells (Appendix Fig S2B).

**Figure 2.**
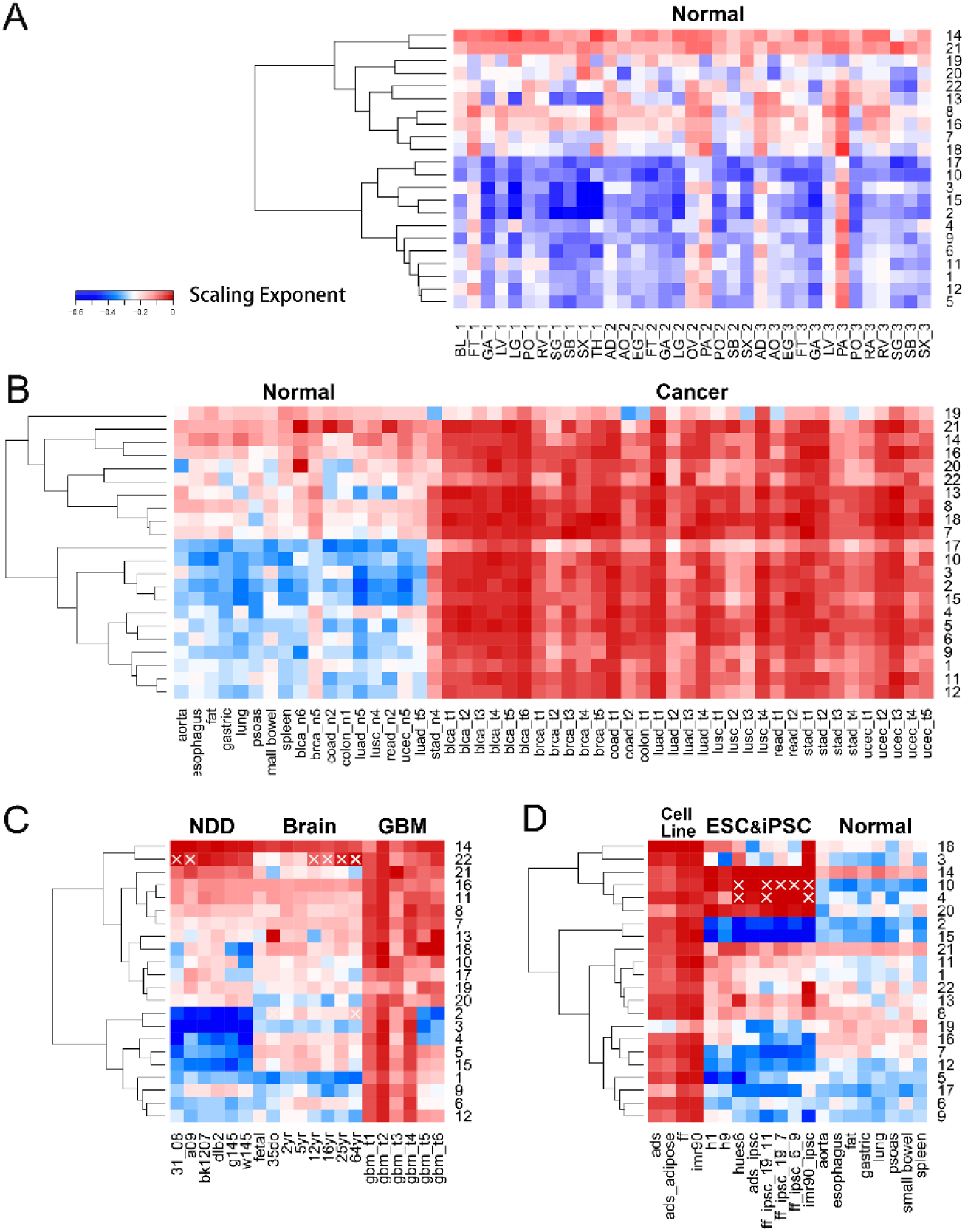
Heatmap clustering for the scaling exponents of all autosomal chromosomes shows differences between different cell classes. Sample labels are summarized in Appendix Data. A, The scaling exponents of normal somatic cells are robust among 3 different individuals and the gland cells segregate from other cells. B, Normal somatic cells (left to right: aorta to ucec_n5) segregate from cancer cells (brca_t1 to ucec_t5). luad_t5 and stad_n4 are further analyzed in Appendix Data. C, Normal brain cells (fetal to 64yr) segregate from glioblastoma (GBM, gbm_t1 to gbm_t6) or neurodegenerative diseases (NDD, 31_08 to w145). Normal somatic cells (aorta to spleen) were shown again for comparison. D, ESCs and iPSCs (h1 to imr90_ipsc) segregate from cell lines including adult stem cell line (ads) and somatic cell lines (ads_adipose, ff, imr90). The chromosomes with correlations deviate from power-law distributions are labeled with white forks (see Appendix Data for more analyses).

In addition, detrended fluctuation analyses (Peng et al, 1992) were also performed which again show the long-range correlation in DNA methylome (Appendix Fig S4). Detrended fluctuation analysis was used previously to describe the long range correlation in DNA sequences (Peng et al, 1992), which is more robust than direct correlation calculation when determining the average behavior of a long range effect.

The significant difference between different cell classes demonstrates that the long-range correlations in DNA methylome cannot be simply originated from DNA sequence (Arneodo et al, 2011; Peng et al, 1992). DNA methylation was previously demonstrated to have long-range correlations by establishing a firm link with A/B compartment (Fortin & Hansen, 2015), suggesting the scale-free property found here for DNA methylation to be originated from chromatin structure, which is discussed later in more detail.

### Clustering on the scaling exponents of chromosomes can be used to discern different cell types

The power law scaling behavior is observed in almost all chromosomes across a large variety of samples (Appendix Fig S2C). Hierarchical clustering for scaling exponents on all autosomal chromosomes demonstrates that most chromosomes behave similarly within each cell class, while chromosome 14 and 21 tend to always have a higher scaling exponent (Fig 2). When all chromosomes are compared, it can be clearly seen that cancer cells are distinguished from normal somatic cells (Fig 2B), consistent with the clustering on cell types (Appendix Fig S3A). Systematic differences are also clearly seen among normal brain cells, glioblastoma (GBM) and neurodegenerative diseases (NDD) (Fig 2C). Different types of neurodegenerative diseases have similar scaling exponents while behaving significantly differently from glioblastoma, possibly highlighting their different pathogenesis (Fig 2C). In addition, the scaling exponent also clearly distinguishes embryonic stem cells (ESCs) and induced pluripotent stem cells (iPSCs) from somatic cell lines and adult stem cell lines (Fig 2D).

When compared to the normal brain cells, all neurodegenerative disease samples analyzed here possess more negative scaling exponents for chromosome 2, 3, 5, and 15, suggesting their common roles associated with the neural diseases. In contrast, chromosome 19 shows little variation among all samples.

A small number of chromosomes possess correlations that deviates from a simple power-law scaling (Appendix Fig S2D). We systematically identify such chromosomes (see Materials and Methods) which are, interestingly, mainly found in certain cells and particular chromosomes, namely the chromosome 22 of the brain samples and chromosomes 4 and 10 of ES cells and iPSCs. Although the atypical power-law behavior of these chromosomes may reflect the large fluctuation of the original methylation data, the clustered behavior could also suggest that these particular chromosomes have peculiar structures and functions which call for further studies. For example, it is known that genes in chromosome 22 are dense and genetic disorder in chromosome 22 is associated with brain abnormalities (McDermid & Morrow, 2002).

### Patchiness of DNA methylation is found along the genome and contributes to the power law scaling

Extensive changes in DNA methylation take place during tumourgenesis (Berman et al, 2012; Hon et al, 2012). In cancer cells, a large amount of long-range DNA hypomethylation was identified, distinct from the DNA methylation of normal cells (Berman et al, 2012). The domain with long-range DNA hypomethylation is termed as partially-methylated-domain (PMD) (Lister et al, 2009). The IMR90 cell line also has such DNA hypomethylation character (Berman et al, 2012; Lister et al, 2009; Lister et al, 2011).

For cancer cells or IMR90 cell line, the whole chromosome can be viewed as composed of alternating low-methylation-level domain (i.e. PMD) and high-methylation-level domain (i.e. non-PMD) in contrast to other cells. That is, the patchiness of DNA methylation for cancer cells or IMR90 cell line is more apparent than normal somatic and stem cells. The scaling exponents for IMR90 and cancer cells are similar to each other (Fig 2). Such a coincidence promoted us to investigate whether the patchiness of DNA methylation contributes to the different scaling exponents of long-range DNA methylation in different cell classes.

Chromosome 1 in IMR90 and h1 cell line are taken for example. There are 34% PMD in IMR90, while h1 lacks PMDs. Namely, IMR90 and h1 have different degrees of DNA methylation patchiness. To understand how such patchiness is generated in IMR90 but not h1 cell, their Hi-C data which are available are used in the next section for structural modeling (Dixon et al, 2012; Rao et al, 2014).

Here we show that the high-low alternative pattern of DNA methylation is enough to mathematically reproduce the slow-decaying correlation in IMR90. we discretized the DNA methylation level of IMR90 and h1 into 1 and 0 with the methylation average as reference value. Specifically, for chromosome 1 of each cell type, we assign a value of 1 to every 200-bp unit with methylation level greater than chromosome average, and 0 to that with methylation level smaller than average. The correlations of the two discrete model series were calculated and shown in Fig 3B. The corresponding correlations of experimental DNA methylation level are also plotted in Fig 3A for comparison. The discrete model series also possess the power law scaling behavior at the kilobase-to-megabase scale (Fig 3B). The comparison between Fig 3A and 3B shows that the discrete model is able to reproduce the different scaling exponents in IMR90 and h1 cell lines, proofing the difference mainly come from the different patchiness of their DNA methylation pattern. That is, the alternation of low and high methylation alternation along the genome in IMR90 results in the lower power law scaling exponent compared to h1 cell line.

**Figure 3.**
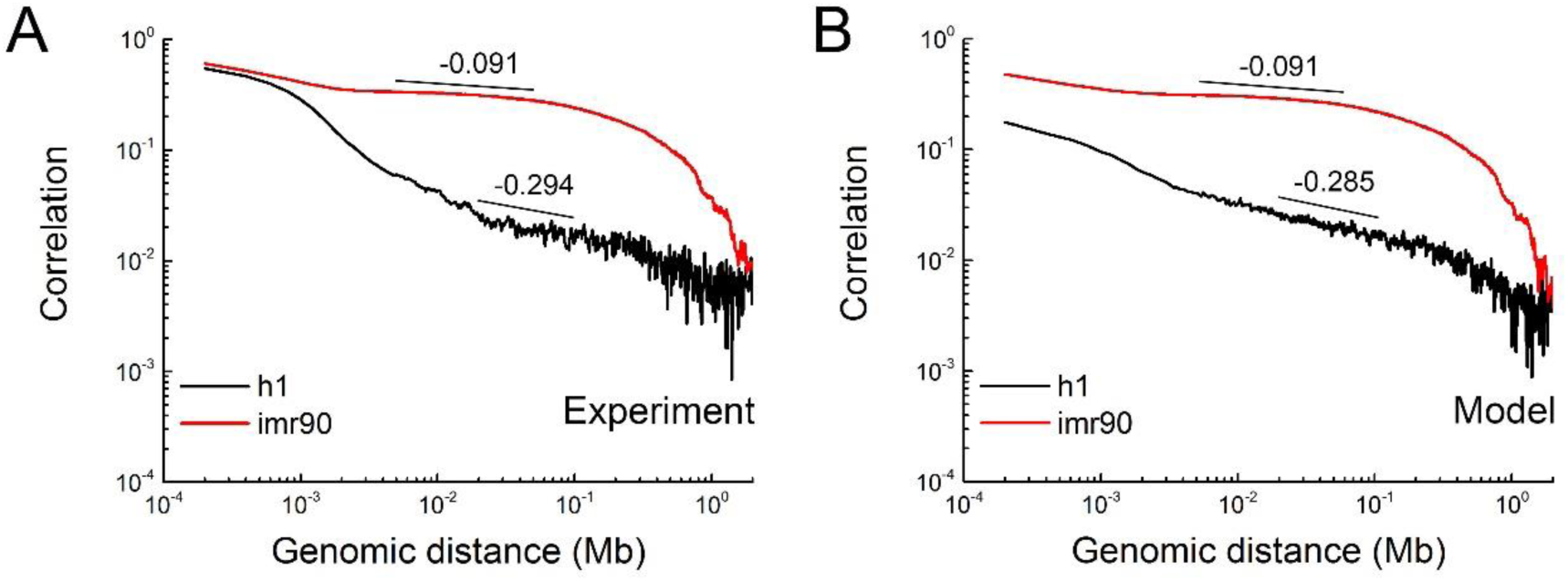
The comparison between the experimental and discrete model DNA methylation correlation indicates that the power law scaling mainly originate from the patchiness of DNA methylation. The long-range correlation of DNA methylation for chromosome 1 in IMR90 and h1 cell line from: A, Experimental methylation data; B, Discrete model.

### Patchiness of DNA methylation is related to chromatin structure

Next we show the patchiness of DNA methylation reflects the packing of DNA in the 3D chromatin structure by mapping methylation level onto the modeled chromatin structure using Hi-C data.

We have developed a polymer modeling strategy using Hi-C data to construct the chromatin structure (see Materials and Methods and preprint: Xie et al, 2016). Hi-C data provide the frequency of physical interactions between any different genomic loci (Lieberman-Aiden et al, 2009), and the frequencies can be further related with spatial distances (Serra et al, 2015). We use structural optimization to obtain the coarse-grained chromatin conformations meeting the distance constraints derived from Hi-C data.

We modeled structures of chromosome 1 from IMR90 and h1, and mapped their DNA methylation levels onto the structures, which are respectively shown in Figs 4A and 4B. The two chromatin structures have obviously different organizations. Chromosome 1 of IMR90 shows a somewhat spherical appearance (Fig 4A), while h1 chromosome adopts a scissor-like conformation (Fig 4B), suggesting structural changes during cellular differentiation. The mapping of DNA methylation level might provide a clue of how the different patchiness of DNA methylation in the two cell lines happen. In IMR90, genomic regions with low methylation levels (colored blue in Fig 4A) segregate from high methylation regions. In contrast, the segregation is not obvious in h1 cell line (Fig 4B).

**Figure 4.**
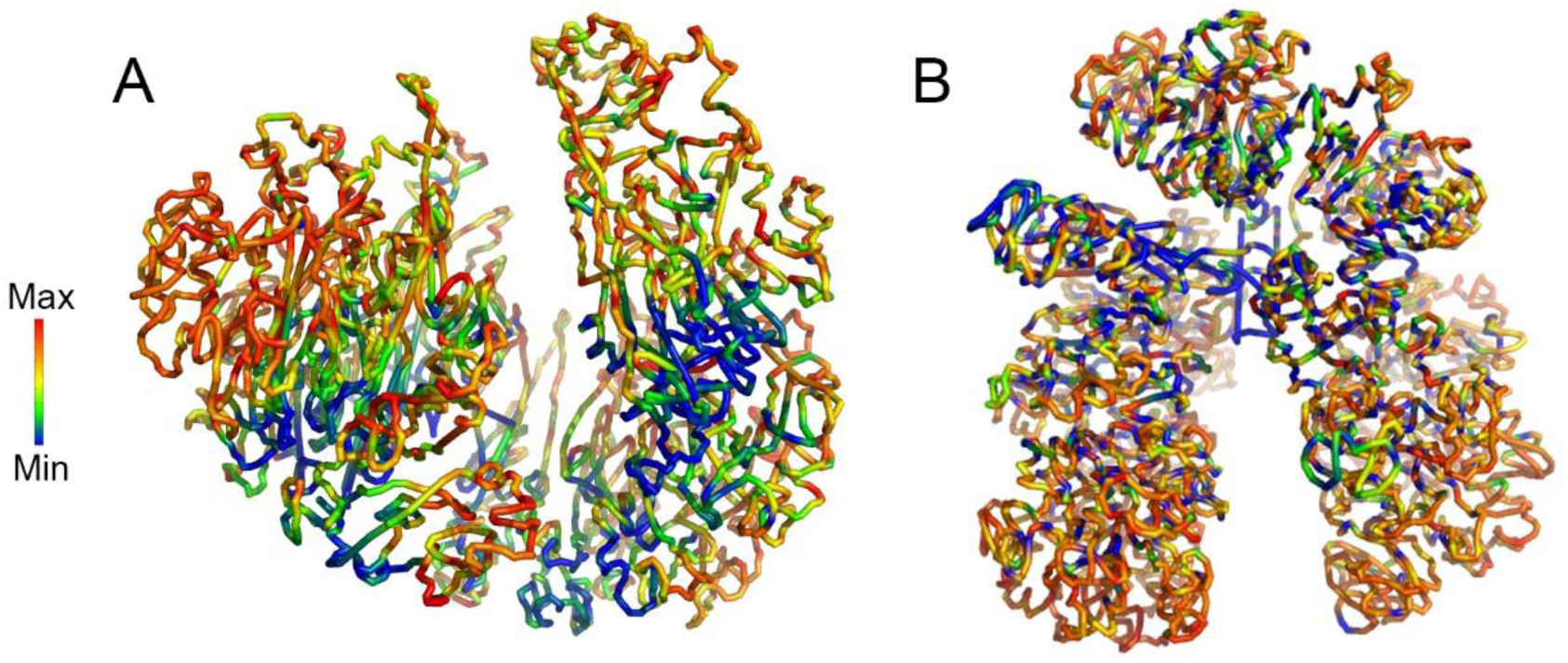
Patchiness of DNA methylation is related to chromatin structure. Modeled chromatin structures of chromosome 1 with mapped methylation level in A, IMR90 and (B) h1. Blue (red) color represents low (high) DNA methylation level.

In our previous work, we have shown that the segregated low methylation regions (PMDs) in IMR90 are related to lamina-associated-domains as well as other genome features (preprint: Xie et al, 2016), showing the origin of different patchiness in IMR90 from h1. These results suggest that the different patchiness of DNA methylation in different cells are related to their different chromatin structures. Thus, the long-range power law correlation for DNA methylation can reflect the spatial organization of chromatin, which in itself is hierarchical.

### Local methylation correlations suggest the different chromatin structure in PMD and non-PMD

In the previous section, we have shown that the long-range correlation of DNA methylation reflects the global packing of DNA in chromatin. Next, we show that the local methylation correlations in PMD and non-PMD reflect their different structures. Breast cancer is used as an example because of the availability of extensive amount of data.

We compared the local correlation of CpG methylation in PMDs and non-PMDs in breast cells (Fig 5A). Consistent with previous studies on IMR90 cell line (Gaidatzis et al, 2014), the decay of PMD correlation clearly shows an obvious periodic behavior at base-resolution, again showing the similarity of the PMD regions in different cells. The non-PMD regions, in contrast, do not show any apparent periodic behavior. The corresponding genomic regions of cancer PMDs in normal cells are defined as PMD-like regions. We found that PMD-like regions have an average methylation level higher than PMD and lower than non-PMD (Fig 6B), and a less obvious methylation correlation periodicity (Fig 5A).

**Figure 5.**
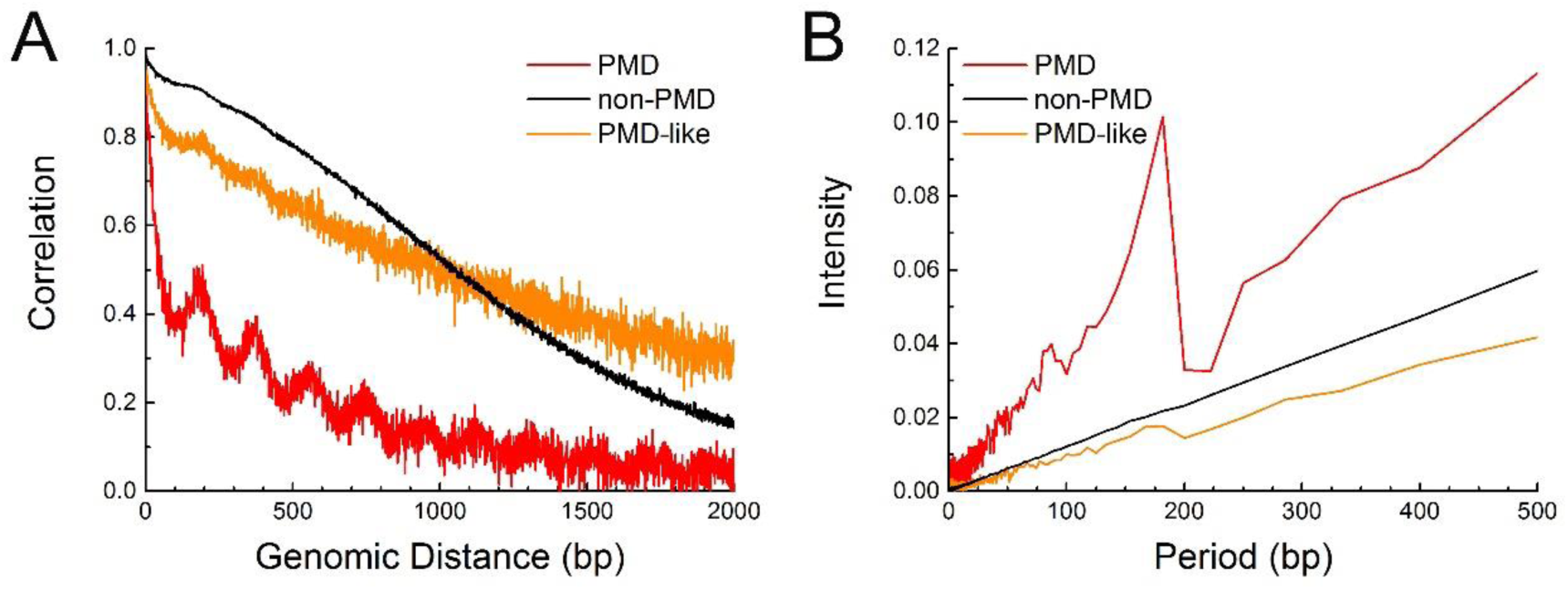
The local correlations of DNA methylation in PMD, non-PMD and PMD-like genomic regions in breast cells. A, Local correlations of DNA methylation. B, FFT is used to deconstruct inherent frequencies of the local correlations.

The periodicities in different genomic regions were then quantified using fast Fourier transform (FFTs) of their local correlations. For PMD, the FFT of its local correlation shows a strong peak at 181 bp. At the similar position, a weaker peak was found for PMD-like regions (Fig 5B). The period of 181 bp is consistent with the nucleosome repeat length, suggesting that this periodicity may come from the regular organization of nucleosomes in PMD. In contrast, the absence of periodicity for non-PMD local correlation indicates the nucleosomes in non-PMD are irregularly spaced.

The regularity of nucleosome arrangement can be weakened by nucleosome depletion or different nucleosome repeat length (NRL). Both the two factors severely affect chromatin organization and compaction. Nucleosome depletion massively influence chromatin flexibility (Arneodo et al, 2011; Diesinger & Heermann, 2009; Ricci et al, 2015). With different nucleosome repeat lengths, the nucleosomes can form 30-nm higher-order chromatin structure or other chromatin fibers (Grigoryev, 2012; Routh et al, 2008). Therefore, the local correlation of DNA methylation suggests that the chromatin structure of PMD and non-PMD is different at kb genomic scale.

Such a conclusion is consistent with our previous analysis of Hi-C data (preprint: Xie et al, 2016). We found the Hi-C pattern for PMD and non-PMD are obviously different which again shows these two domains have different spatial organization. From Hi-C data, it is easy to see that all the PMDs have uniform physical contact within its interior, while the majority of non-PMD contains localized interaction domains.

### Gene expression in PMDs and PMD-like regions are repressed

To understand how the patchiness of DNA methylation is related to biological functions, we analyze the gene expression in PMDs and non-PMDs. As explained in Materials and Methods, we analyzed the 4 tumor-normal sample pairs in TCGA. It was previously found that the PMDs in IMR90 correlate with repressive and anti-correlate with active histone marks (Berman et al, 2012). Consistent with earlier studies (Lister et al, 2011; Schroeder et al, 2013; Schultz et al, 2015), we find that genes within PMDs in cancer samples tend to be transcriptionally repressed (Appendix Tables S6 and S7, Appendix Fig S5) and, interestingly, these genes are related to specific functions. Genes within cancer PMDs mainly relate to GO terms such as cell membrane, glycoprotein, disulfide bond, olfaction, cadherin and receptor (Appendix Table S8), which suggests that some intra-PMD genes regulating cell communication tend to be repressed. Besides, almost all housekeeping genes (3794 of 3796) are located outside PMDs, consistent with their essential role in the fundamental cellular function (Appendix Table S8).

Taking the brca_t5 tumor sample as an example, there are 473 genes intersecting with PMDs, of which 305 are located within the PMD body and especially, 156 are in the PMD center (defined as the central 60% of PMD), indicating that most genes embed in the PMD body. Besides, among the 473 genes intersecting with PMDs, 57.7% of them have non-CGI promoters (the definition of non-CGI and CGI promoter is from (Saxonov et al, 2006)) and this ratio is significantly higher than that of all genes (34.2%, 8420 of total 24630 genes) which indicates that genes with non-CGI promoters are enriched in PMDs (Appendix Table S8).

Besides the repressed gene expression level, we also find the repression degree correlates with the PMDs length in the 4 tumor-normal sample pairs. Fig 6A shows the correlation between gene repression levels and PMD lengths in brca and the results in other 3 tumor-normal sample pairs (coad, luad, ucec) are presented in Appendix Fig S6. With the increasing length of PMDs or PMD-like regions, the percentage of repressed gene increases, which may result from these genomic regions being buried in the compact chromatin regions in 3D space and probably also impedes the binding with TFs, RNA polymerase or other regulators. The gene density of PMDs (2.737 in brca_t5 sample) is also much lower than that of non-PMDs (6.526 in brca_t5 sample), indicating the gene sparsity in PMDs.

**Figure 6.**
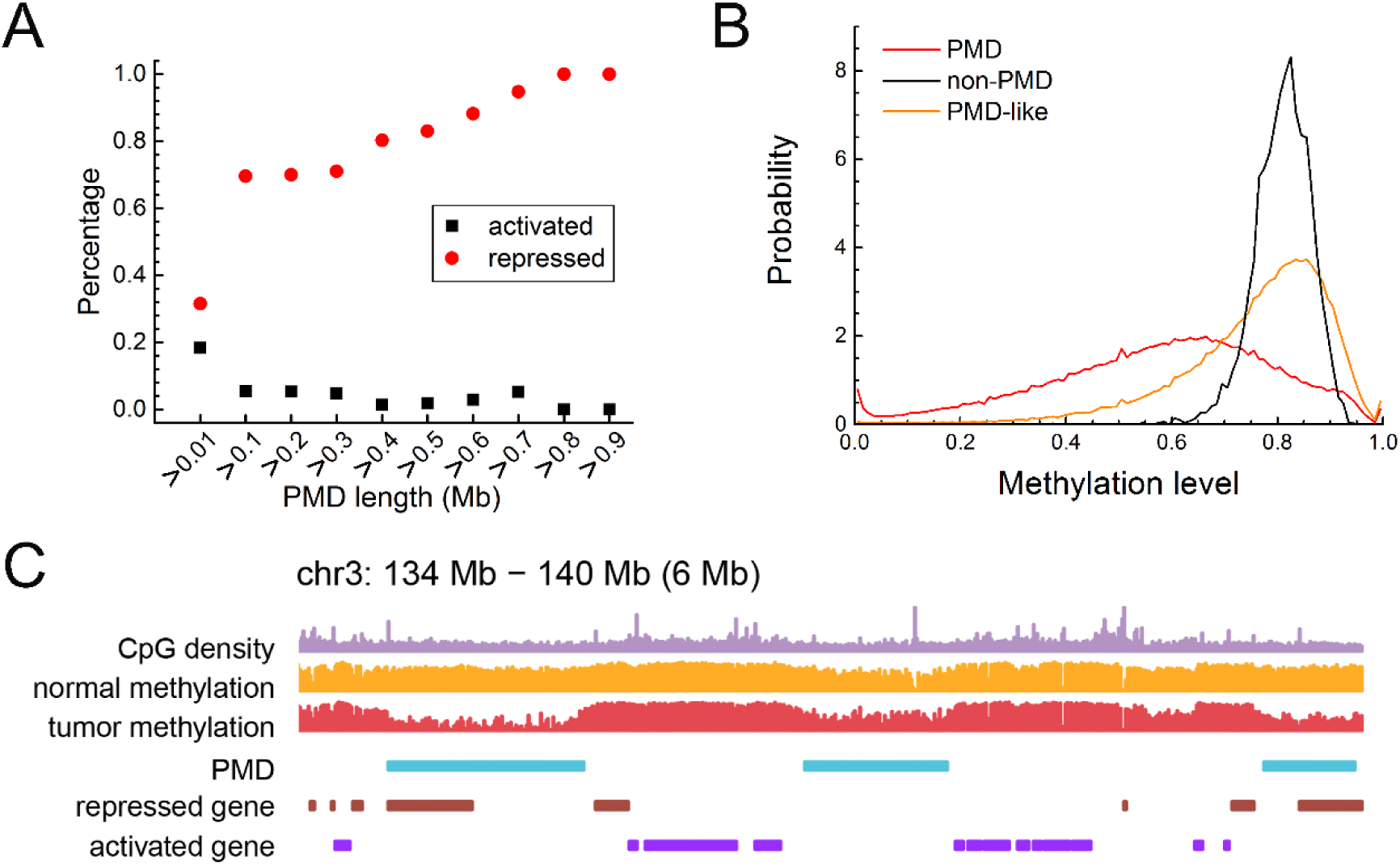
Gene expression and sequence property of PMD and non-PMD. A, Percentage of genes that are transcriptionally repressed or activated in oncogenesis as a function of PMD length in brca_t5-brca_n5 sample pair. The comparisons in coad_t2-coad_n2 and ucec_t5-ucec_n5 sample pairs are shown in Appendix Fig S6. B, Distribution of methylation level of PMD, non-PMD and PMD-like regions in brca samples. C, Plot of a representative region showing the relation among CpG density, methylation levels before and after oncogenesis, PMD, and repressed and activated genes in oncogenesis. In A, B and C, the data of brca were used.

Furthermore, the average expression level of genes in PMD-like regions is lower than that in non-PMDs and higher than that in PMDs which is consistent with PMD-like regions’ intermediate behavior in the DNA methylation level (Fig 6B) and the periodicity of local methylation correlation (Fig 5). The intermediate properties and the similar genomic locations of PMD-like regions to the PMDs imply the role of PMD-like regions in the tumor development. PMD-like regions may be the precursor of tumor PMDs in which the genes regulating cell communications are further repressed. Interestingly, PMD or PMD-like domains tend to lie in genomic regions with lower CpG density (Fig 6C), suggesting their belonging to different isochores (Costantini et al, 2006). In this analysis, the Fisher’s exact test between the expression level of PMDs and PMD-like regions and that between PMDs and non-PMDs is shown in Appendix Table S7. The average expression level in PMDs, PMD-like regions and non-PMDs is shown in Appendix Table S6. The comparison of expression level in PMD and PMD-like regions in the 4 tumor-normal sample pairs is shown in Appendix Fig S6.

## Discussion

### Power law scaling in cancer cells is not caused by the lower average methylation level or copy number variation

One difference between cancer and normal somatic cells in DNA methylation is that the former appears to be demethylated in PMDs compared to the latter. To show that the more sustained correlation of cancer cell DNA methylation is not caused by this overall demethylation, we checked the scaling exponents of methylation correlations among cells with large variations in methylation levels. We calculated the methylation correlations of human inner cell mass (ICM) (Guo et al, 2014) and primordial germ cells (PGCs) (Guo et al, 2015). The average methylation level of these cells are both significantly lower than normal as well as cancer cells (Appendix Fig S7A). However, the scaling exponents for ICM and PGC are nearly the same as normal cells and much lower than cancer cells (Appendix Fig S7B). Thus it can be concluded that average methylation level does not determine the different scaling exponents across cell classes.

Since tumor samples are enriched in copy number variations (CNVs), we showed that the DNA methylation correlation is affected little by CNVs in cancer cells. For example, the long-range methylation correlation for chromosome 1 behaves similarly whether the CpG sites within CNVs of brca_t5 tumor sample are included or excluded (Appendix Fig S7C) in the correlation calculations.

Through exploiting the chromatin structure modeled based on Hi-C data and the underlying long-range and local correlations of DNA methylome, our study provides a comprehensive view of the flow of genetic information, connecting DNA sequence, CpG methylation, local and long-range chromatin structure, and gene expression. In normal somatic cells, DNA sequences with low CpG density correlate with low methylation levels and low expression levels (PMD-like). The development of cancers is associated with further decrease of the average methylation level in PMD-like regions, some of which turn into PMDs containing further suppressed genes. The correlation of methylation shows consistent differences among different classes of cells, including normal somatic cells, cancer cells, brain cells, gland cells and stem cells, but are highly conserved within each class. The clear cell class dependence of the long-range power law scaling in methylation correlation shows that it can serve as a simple measure to discriminate cells at normal and pathological states. Such a finding points to a new direction in the analysis of development of different diseases, for example, cancers and neurodegenerative diseases at the chromatin level.

## Materials and Methods

### Sources of whole-genome bisulfite sequencing data

In this work, we used the whole genome bisulfite sequencing (WGBS) data for different cells, including 36 somatic cells, 42 cancer cells and the corresponding normal cells, 8 human brain cells, 1 mouse brain cells, 12 stem cells and related cells, 6 cells with neurodegenerative diseases (105 in total). All the methylomes were summarized in Appendix Data, Appendix Tables S1-S5, including the references, URLs, sample details. The methylomes of cancer samples were downloaded from the The Cancer Genome Atlas (TCGA) project. Of all the samples in TCGA, 9 types of cancer samples have WGBS data and these 9 samples were used.

Hg18 reference genome was used for human brain cells and human ESCs. The other cells used Hg19 as reference genome. We examined that reference genome has little effect on the methylation correlation found here (Appendix Fig S7D).

### Identification of PMDs, non-PMDs and PMD-like regions

PMDs were identified genome-wide in tumor samples using a sliding window approach and the parameters in (Lister et al, 2009) were adopted in this work. The window size was set as 10 kb. A region was identified as a PMD if there were at least 10 methylated (beta value greater than 0) CpG dinucleotide within, of which the average methylation level was less than 0.7. The contiguous PMD windows were then merged into a longer PMD. Only PMDs with lengths longer than 100 kb are used in the following analysis. Non-PMDs are identified as the complementary set of the PMDs and only non-PMDs whose lengths are greater than 100 kb are used. PMD-like regions were defined in non-cancer samples as the corresponding genomic regions of cancer PMDs. Thus, PMD-like regions are defined only for the non-cancer samples which have corresponding cancer samples.

### Calculation of scaling exponents

The scaling exponents of the long-range power law correlations were calculated as the maximum slope of the fitted double-log correlation data in the genomic region of kilobase to megabase. To systematically identify those chromosomes whose slope of double-log plot of methylation correlation is not well-defined in the concerned (kilobase to megabase) region, we calculated the standard deviation of the first order derivatives for each fitted double-log plot. Low standard deviations indicate linear behavior with small slope fluctuation for methylation correlation, while high standard deviations indicate large fluctuation.

### Fast Fourier Transform of the local correlation

Fast Fourier transform was performed on the local correlation of the 3 genomic regions (PMDs, PMD-like regions and non-PMDs) respectively. To avoid the finite length effect and influences of length distribution of genomic regions, we chose PMDs with a genomic length of 1 to 2 Mb from all breast cancer samples, corresponding PMD-like regions from the breast normal sample, and non-PMDs with a genomic length of 1 to 2 Mb from the brca_t1 sample to calculate PMD, PMD-like and non-PMD Fourier transform, respectively.

### Gene expression analysis

The level3 RSEM data from TCGA RNAseq version2 was downloaded from https://portal.gdc.cancer.gov. The RSEM data were then converted to Transcripts Per Million (TPM) by multiplying by 10^6^. To compare the differentially expressed genes between tumor and normal samples, we chose the tumor-normal sample pairs that were taken from the same patient and gene expression data of both tumor and normal samples were available. We finally obtained 4 tumor-normal pairs, namely, brca_t5-brca_n5, coad_t2-coad_n2, luad_t5-luad_n5 and ucec_t5-ucec_n5. We compared gene expression for these 4 tumor-normal pairs.

Genes with intragenic regions intersecting with tumor PMDs (or PMD-like regions in corresponding normal samples) were identified. Then gene expression change was calculated as TPM_*tumor*_ /TPM_*normal*_ for each gene. These genes were divided into four categories: activated genes (fold change ≥ 2), repressed genes (fold change ≤ 0.5), specifically expressed in tumor sample(TPM_*normal*_ = 0 and TPM_*tumor*_ ≠ 0) and specifically expressed in normal sample(TPM_*normal*_ ≠ 0 and TPM_*tumor*_ = 0). We also defined the gene density of the genome and the specific genomic regions like PMDs as the number of genes per million base pairs. Gene functional classification was carried out using DAVID (The Database for Annotation, Visualization and Integrated Discovery)(Huang et al, 2009a) (Huang et al, 2009b). Housekeeping genes list is downloaded from https://www.tau.ac.il/∼elieis/HKG/ (Eisenberg & Levanon, 2013).

### Structural modeling using Hi-C data

We developed a restraint-based method to construct an ensemble of 3D chromosome models (preprint: Xie et al, 2016). The method was verified by the reproduction of experimental Hi-C contact frequencies. In our method, chromosome was coarse-grained as a polymer chain consisting of a string of beads. The Hi-C data for IMR90 and h1 cell line were obtained from Rao et al (Rao et al, 2014) and Dixon et al (Dixon et al, 2012), respectively. According to the resolution of Hi-C data, in our modeling for IMR90 and h1, each bead respectively represents a 50 kb or 40 kb genomic region. The polymer structure was optimized according to distance restraints derived from Hi-C data. To achieve this, we first converted the contact frequency matrix measured by the Hi-C experiment to a distance matrix which provides the spatial restraints for the coarse-grained beads. Then we performed molecular dynamics simulations starting from randomly generated initial conformations using biased potentials to generate an ensemble of conformations based on the restraint distance matrix. Further modeling details and validation were presented in Xie *et al* (preprint: Xie et al, 2016).

## Acknowledgements

We thank Fuchou Tang, Chengqi Yi and Xiaoliang S. Xie for helpful comments on the manuscript. The results shown here are partly based upon data generated by the TCGA Research Network: http://cancergenome.nih.gov/. We are grateful to Manel Esteller for providing the methylome of neurodegenerative diseases. This work was supported by National Natural Science Foundation of China [21573006, 21233002 and 91427304 to Y.Q.G.].

## Author contributions

Y.Q.G designed the research. L.Z., W.J.X., S.L. and L.M. performed research. W.J.X., L.Z., S.L. and L.M. wrote the manuscript. G.C. contributed to the analysis of the data.

## Conflict of interest statement

The authors declare that they have no conflict of interest.

## References

Aranda S, Mas G, Di Croce L (2015) Regulation of gene transcription by Polycomb proteins. Sci Adv 1: e1500737

Arneodo A, Vaillant C, Audit B, Argoul F, d’Aubenton-Carafa Y, Thermes C (2011) Multi-scale coding of genomic information: From DNA sequence to genome structure and function. Phys Rep 498: 45–188

Bell AC, Felsenfeld G (2000) Methylation of a CTCF-dependent boundary controls imprinted expression of the Igf2 gene. Nature 405: 482–485

Berman BP, Weisenberger DJ, Aman JF, Hinoue T, Ramjan Z, Liu Y, Noushmehr H, Lange CP, van Dijk CM, Tollenaar RA, Van Den Berg D, Laird PW (2012) Regions of focal DNA hypermethylation and long-range hypomethylation in colorectal cancer coincide with nuclear lamina-associated domains. Nat Genet 44: 40–46

Boettiger AN, Bintu B, Moffitt JR, Wang S, Beliveau BJ, Fudenberg G, Imakaev M, Mirny LA, Wu CT, Zhuang X (2016) Super-resolution imaging reveals distinct chromatin folding for different epigenetic states. Nature 529: 418–422

Cedar H, Bergman Y (2009) Linking DNA methylation and histone modification: patterns and paradigms. Nat Rev Genet 10: 295–304

Chodavarapu RK, Feng S, Bernatavichute YV, Chen P-Y, Stroud H, Yu Y, Hetzel JA, Kuo F, Kim J, Cokus SJ, Casero D, Bernal M, Huijser P, Clark AT, Kramer U, Merchant SS, Zhang X, Jacobsen SE, Pellegrini M (2010) Relationship between nucleosome positioning and DNA methylation. Nature 466: 388–392

Clauset A, Shalizi CR, Newman MEJ (2009) Power-law distributions in empirical data. Siam Rev 51: 661–703

Costantini M, Clay O, Auletta F, Bernardi G (2006) An isochore map of human chromosomes. Genome Res 16: 536–541

Dekker J (2008) Gene regulation in the third dimension. Science 319: 1793–1794

Diesinger PM, Heermann DW (2009) Depletion effects massively change chromatin properties and influence genome folding. Biophys J. 97: 2146–2153

Dixon JR, Selvaraj S, Yue F, Kim A, Li Y, Shen Y, Hu M, Liu JS, Ren B (2012) Topological domains in mammalian genomes identified by analysis of chromatin interactions. Nature 485: 376–380

Eisenberg E, Levanon EY (2013) Human housekeeping genes, revisited. Trends Genet 29: 569–574

Eugene VK, Yuri IW, Georgy PK (2006) Power Laws, Scale-Free Networks and Genome Biology, New York: Springer.

Fortin JP, Hansen KD (2015) Reconstructing A/B compartments as revealed by Hi-C using long-range correlations in epigenetic data. Genome Biol 16: 180

Gaidatzis D, Burger L, Murr R, Lerch A, Dessus-Babus S, Schubeler D, Stadler MB (2014) DNA sequence explains seemingly disordered methylation levels in partially methylated domains of Mammalian genomes. PLOS Genet 10: e1004143

Galupa R, Heard E (2015) X-chromosome inactivation: new insights into cis and trans regulation. Curr Opin Genet Dev 31: 57–66

Gibcus JH, Dekker J (2013) The hierarchy of the 3D genome. Mol Cell 49: 773–782

Grigoryev SA (2012) Nucleosome spacing and chromatin higher-order folding. Nucleus 3: 493–499

Guo F, Yan LY, Guo HS, Li L, Hu BQ, Zhao YY, Yong J, Hu YQ, Wang XY, Wei Y, Wang W, Li R, Yan J, Zhi X, Zhang Y, Jin HY, Zhang WX, Hou Y, Zhu P, Li JY et al (2015) The transcriptome and DNA methylome landscapes of human primordial germ cells. Cell 161: 1437–1452

Guo HS, Zhu P, Yan LY, Li R, Hu BQ, Lian Y, Yan J, Ren XL, Lin SL, Li JS, Jin XH, Shi XD, Liu P, Wang XY, Wang W, Wei Y, Li XL, Guo F, Wu XL, Fan XY et al (2014) The DNA methylation landscape of human early embryos. Nature 511: 606–610

Hon GC, Hawkins RD, Caballero OL, Lo C, Lister R, Pelizzola M, Valsesia A, Ye Z, Kuan S, Edsall LE, Camargo AA, Stevenson BJ, Ecker JR, Bafna V, Strausberg RL, Simpson AJ, Ren B (2012) Global DNA hypomethylation coupled to repressive chromatin domain formation and gene silencing in breast cancer. Genome Res 22: 246–258

Huang DW, Sherman BT, Lempicki RA (2009a) Bioinformatics enrichment tools: paths toward the comprehensive functional analysis of large gene lists. Nucleic Acids Res 37: 1–13

Huang DW, Sherman BT, Lempicki RA (2009b) Systematic and integrative analysis of large gene lists using DAVID bioinformatics resources. Nat Protoc 4: 44–57

Jenuwein T, Allis CD (2001) Translating the histone code. Science 293: 1074–1080

Levine M, Cattoglio C, Tjian R (2014) Looping back to leap forward: transcription enters a new era. Cell 157: 13–25

Lieberman-Aiden E, van Berkum NL, Williams L, Imakaev M, Ragoczy T, Telling A, Amit I, Lajoie BR, Sabo PJ, Dorschner MO, Sandstrom R, Bernstein B, Bender MA, Groudine M, Gnirke A, Stamatoyannopoulos J, Mirny LA, Lander ES, Dekker J (2009) Comprehensive mapping of long-range interactions reveals folding principles of the human genome. Science 326: 289–293

Lister R, Pelizzola M, Dowen RH, Hawkins RD, Hon G, Tonti-Filippini J, Nery JR, Lee L, Ye Z, Ngo QM, Edsall L, Antosiewicz-Bourget J, Stewart R, Ruotti V, Millar AH, Thomson JA, Ren B, Ecker JR (2009) Human DNA methylomes at base resolution show widespread epigenomic differences. Nature 462: 315–322

Lister R, Pelizzola M, Kida YS, Hawkins RD, Nery JR, Hon G, Antosiewicz-Bourget J, O’Malley R, Castanon R, Klugman S, Downes M, Yu R, Stewart R, Ren B, Thomson JA, Evans RM, Ecker JR (2011) Hotspots of aberrant epigenomic reprogramming in human induced pluripotent stem cells. Nature 471: 68–73

McDermid HE, Morrow BE (2002) Genomic disorders on 22q11. Am J Hum Genet 70: 1077–1088

Naumova N, Imakaev M, Fudenberg G, Zhan Y, Lajoie BR, Mirny LA, Dekker J (2013) Organization of the mitotic chromosome. Science 342: 948–953

Peng CK, Buldyrev SV, Goldberger AL, Havlin S, Sciortino F, Simons M, Stanley HE (1992) Long-range correlations in nucleotide sequences. Nature 356: 168–170

Pombo A, Dillon N (2015) Three-dimensional genome architecture: players and mechanisms. Nat Rev Mol Cell Biol 16: 245–257

Rao SS, Huntley MH, Durand NC, Stamenova EK, Bochkov ID, Robinson JT, Sanborn AL, Machol I, Omer AD, Lander ES, Aiden EL (2014) A 3D map of the human genome at kilobase resolution reveals principles of chromatin looping. Cell 159: 1665–1680

Ricci MA, Manzo C, Garcia-Parajo MF, Lakadamyali M, Cosma MP (2015) Chromatin fibers are formed by heterogeneous groups of nucleosomes in vivo. Cell 160: 1145–1158

Routh A, Sandin S, Rhodes D (2008) Nucleosome repeat length and linker histone stoichiometry determine chromatin fiber structure. Proc Natl Acad Sci U S A 105: 8872–8877

Saxonov S, Berg P, Brutlag DL (2006) A genome-wide analysis of CpG dinucleotides in the human genome distinguishes two distinct classes of promoters. Proc Natl Acad Sci U S A 103: 1412–1417

Schneider R, Grosschedl R (2007) Dynamics and interplay of nuclear architecture, genome organization, and gene expression. Gene Dev 21: 3027–3043

Schroeder DI, Blair JD, Lott P, Yu HO, Hong D, Crary F, Ashwood P, Walker C, Korf I, Robinson WP, LaSalle JM (2013) The human placenta methylome. Proc Natl Acad Sci U S A 110: 6037–6042

Schultz MD, He Y, Whitaker JW, Hariharan M, Mukamel EA, Leung D, Rajagopal N, Nery JR, Urich MA, Chen H, Lin S, Lin Y, Jung I, Schmitt AD, Selvaraj S, Ren B, Sejnowski TJ, Wang W, Ecker JR (2015) Human body epigenome maps reveal noncanonical DNA methylation variation. Nature 523: 212–216

Serra F, Di Stefano M, Spill YG, Cuartero Y, Goodstadt M, Bau D, Marti-Renom MA (2015) Restraint-based three-dimensional modeling of genomes and genomic domains. FEBS Lett 589: 2987–2995

Xie WJ, Meng L, Liu S, Zhang L, Cai X, Gao YQ, (2016), Structural modeling of chromatin integrates genome features and reveals chromosome folding principle. bioRxiv doi: http://dx.doi.org/10.1101/085167 (To appear in Scientific Reports).

